# Intracerebral recordings in humans reveal gradual emergence of musical beat representation across the dorsal auditory pathway

**DOI:** 10.64898/2026.05.22.727121

**Authors:** T Lenc, J Jonas, S Colnat-Coulbois, B Rossion, S Nozaradan

## Abstract

When experiencing music, humans readily perceive and move along with a periodic beat. This ability has been proposed to rely on an enhanced representation of the beat periodicity in brain activity. However, whether this beat representation is achieved in sensory areas or whether it involves associative brain regions remains debated. Here, we addressed this question using intracerebral depth electrodes implanted in 13 human individuals to record local field potentials directly inside the brain. Participants were presented with an acoustic rhythm known to induce perception of a consistent beat across Western adults. The rhythmic stimulus elicited significant responses in a number of areas, especially superior temporal, parietal and frontal cortices located along the dorsal auditory pathway. Notably, these regions, including the primary auditory cortex but also frontal motor regions, showed significantly enhanced beat periodicity as compared to a biomimetic model of subcortical auditory processing. This beat representation was further sharpened in the inferior parietal lobe, indicating that this associative region may play a key role in the mapping between rhythmic inputs and perceptual templates of beat, in line with its posited function as a sensory-motor interface. Together, these findings provide direct evidence for a gradual transformation of rhythmic sensory input into an abstract representation of beat periodicity. This process appears to rely on the dorsal auditory pathway as a functional network supporting the categorization of rhythmic stimuli into behaviorally relevant timing templates that may be experienced as the beat.

## Introduction

Humans possess an outstanding capacity to coordinate movement in time with both robustness and flexibility [1]. This is well illustrated in collective practices such as music. In these practices, coordinated behaviors imply internal representations of time that are shared across individuals [2]. These internal representations often take the form of the beat, that is, regularly recurring pulses which are elicited yet do not necessarily correspond one-to-one to the temporal structure of sensory inputs [3–5]. Specifically, the same beat can be induced by a wide range of rhythmic inputs whose temporal structure may differ substantially. In other words, beat perception can be conceived as a form of perceptual categorization - a fundamental brain function that allows reproducible selection of a given internal template (here, a beat) across a wide range of physically distinct sensory inputs [6,7].

Evidence for high-level beat categorization comes from a large body of work showing that humans can generate movements aligned with a periodic beat regardless of whether it is explicitly cued by periodic acoustic landmarks [8–14]. In line with behavioral work, there is abundant evidence that the human brain selectively amplifies the representation of beat periodicity when it is not prominent in the low-level features of the sensory input, as captured with non-invasive surface electroencephalography (EEG) [8,9,12–19]. However, the *functional brain network* supporting this transformation is still largely unknown.

To address this issue, recent work used intracerebral EEG by means of depth-electrodes implanted within the brain of patients for the treatment of refractory epilepsy. This work showed a significant enhancement of the beat periodicity in the human temporal lobe, including in the Heschl’s gyrus, which is commonly taken as an anatomical landmark of the primary auditory cortex [20]. What remains unclear is whether this periodized representation is further gradually transformed into an abstract, perceptually relevant format in higher-order associative regions, particularly along the dorsal auditory pathway. Indeed, associative areas, especially regions involved in motor planning and control, have been widely posited as core components of the functional network supporting beat perception across several influential theoretical frameworks [5,21–25]. Empirical support for this notion has primarily come from two main lines of research.

On the one hand, functional magnetic resonance imaging (fMRI) studies have repeatedly shown differential activation of frontal regions, including ventral, dorsal, and medial premotor cortex, in response to rhythmic stimuli that elicit the perception of a periodic beat [26–28]. However, due to its limited temporal resolution and its emphasis on overall activation levels, fMRI provides only indirect insight into how rhythmic sensory inputs are represented in a relative or categorical manner, and how such representations are transformed to support behavior.

On the other hand, electrophysiological recordings in macaque monkeys have shown that neuronal populations in medial premotor cortex exhibit periodic trajectories that reflect internal representations of beat-based timing [29–31]. However, to date, such dynamic trajectories have been observed only in highly trained monkeys performing overt periodic movements or a temporal expectation task, and only in response to strictly periodic sensory inputs such as metronome. Moreover, compared with humans, non-human primates appear to show a limited ability to form abstract beat representations when beat periodicity is not marked by salient acoustic landmarks [6,32, but see 33]. Consequently, the extent to which these findings inform on the neural basis of human musical beat perception remains uncertain.

Here, we go beyond the above-mentioned issues in several critical ways. First, we use intracerebral EEG to measure human brain activity with high temporal and spatial resolution across auditory cortex as well as several higher-order associative regions previously proposed to play a role in beat processing. The excellent temporal resolution of this approach allows us to build on a robust and well-validated analytical framework [6,7,34,35] to quantify and compare the relative prominence of beat representations in local field potentials recorded across brain regions. At the same time, the high spatial resolution of this approach provides a particularly detailed view of how beat-related representations are distributed across auditory and associative cortices. Second, we use a rhythmic stimulus that lacks periodic acoustic landmarks, allowing us to dissociate between the encoding of acoustic features and the representation of a periodic beat. Third, we record brain activity elicited during listening rather than active movement, allowing us to probe automatic perceptual categorization independently of motor execution and task demands. Fourth, we directly compare the obtained neural responses to a biologically plausible model of brainstem auditory processing. This allows us to quantify the emergence of a sharpened representation of the beat in the intracerebral data beyond subcortical auditory processing.

We expected to find gradual enhancement of beat periodicity along the dorsal auditory pathway, with progressive transformation from (i) encoding of stimulus features in the auditory brainstem, to (ii) above chance enhancement of beat periodicity in the auditory cortex, and finally to (iii) more abstract representation of beat periodicity in higher-order associative regions, with sharpened representation of the beat and, inversely, reduced representation of the acoustic features. Such a gradient would indicate a pivotal role of higher-level mechanisms in mapping between rhythmic input and the perceived beat, consistent with prominent theoretical accounts [5,21–25].

Alternatively, beat periodicity may become prominent already within the temporal lobe auditory cortex, without further enhancements in higher-order associative regions. Such an outcome would align with the view that the superior temporal lobe is capable of generating abstract, higher-level representations to support perception [36–41]. This result would also open to non-hierarchical models of functional connections between sensory and higher-order associative cortices, consistent with recent intracerebral studies in other domains such as speech [42] and face recognition [43].

## Results

We recorded intracerebral EEG (also referred to as stereo-encephalography) in 13 participants as they listened to a seamlessly looped rhythmic pattern while remaining still (Fig 1A). This rhythm has been extensively used in previous work and shown to elicit perception of a beat with consistent period across Western adults [8,9,13,14]. Critically, the temporal structure formed by individual sounds making up the rhythm did not match the typically perceived periodic beat. In other words, the “weakly-periodic” structure of the rhythm allowed us to tease apart the neural representation of acoustic features and of the periodic beat.

**Fig 1.**
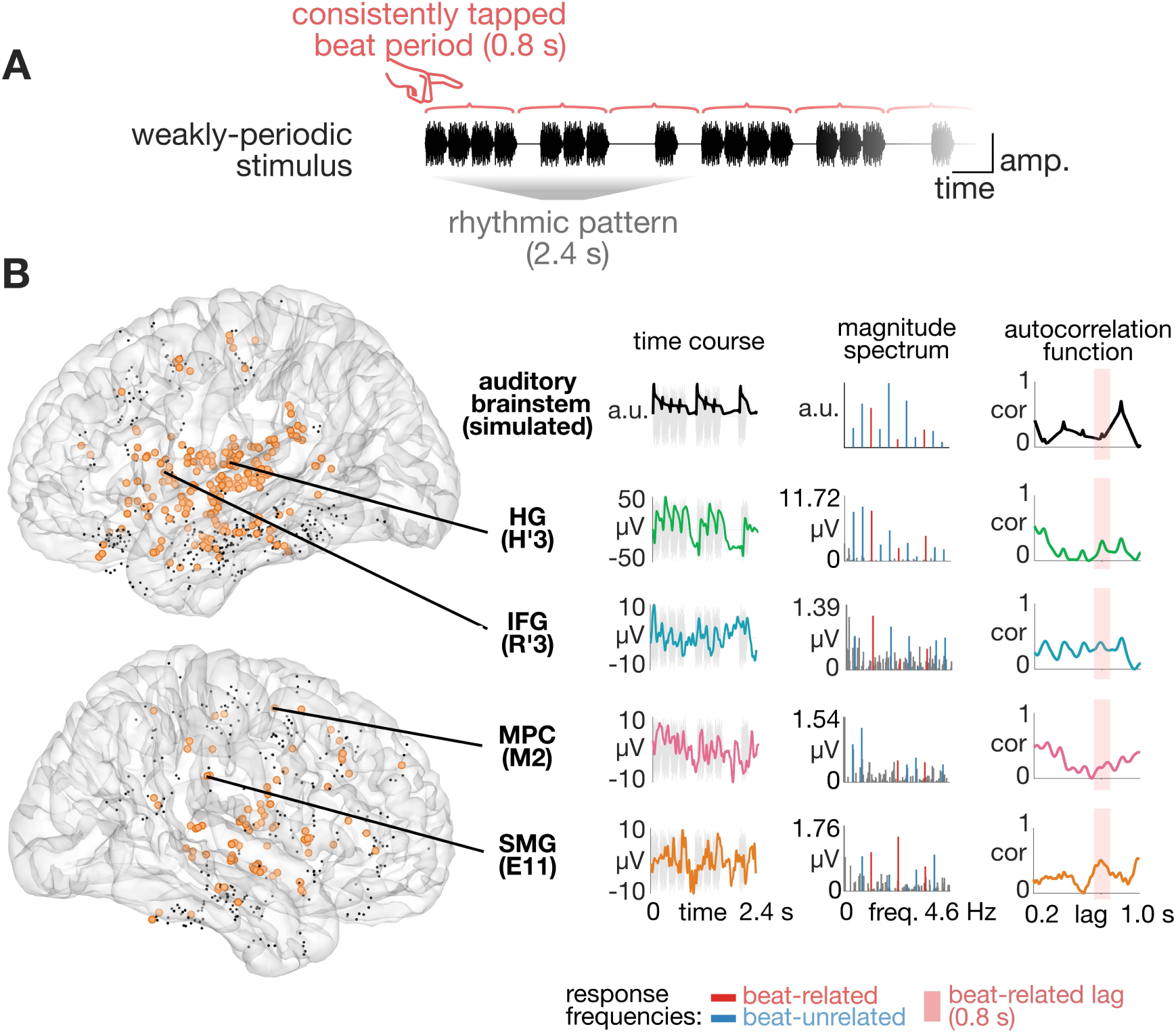
Auditory stimulus and electrode coverage. (A) Segment of the stimulus sequence (audio waveform in black) based on the repeating weakly-periodic rhythmic pattern (indicated in gray below). The beat period consistently tapped by Western adults in response to this stimulus [8,9,13,14,61] is indicated on the top with red brackets. (B) Intracerebral EEG electrode coverage is shown on an MNI brain template. Each point represents an individual recording contact. Responsive contacts are shown as orange circles. Nonresponsive contacts are shown as black dots. Example contacts from regions of interested relevant for the current study are highlighted with arrows. For each example contact (and for the simulated auditory brainstem response), insets show the average response time course to one repetition of the rhythmic pattern (left), magnitude spectrum (middle) and autocorrelation function (right). Frequencies related to the beat periodicity are shown in red. Beat-unrelated frequencies used for magnitude normalization (see Methods) are shown in blue. Autocorrelation lag corresponding to the consistently perceived beat period is indicated by a pink vertical bar.

To demonstrate this, we used a biologically-plausible model simulating responses to the stimulus sequence from neuronal populations of the inferior colliculus in the auditory brainstem [44–46]. From this simulated brainstem response, we measured the relative prominence of the beat periodicity, hereafter “beat index”. This index quantifies how periodic (i.e., recurrent) the response is at the rate of the beat by capturing how much the beat-related frequencies (harmonics of the beat period) stand out in the magnitude spectrum of the signal (values of -1 and 1 indicating minimally and maximally prominent beat periodicity respectively; see Methods for further details).

As shown in Fig 1B, the simulated brainstem response shows a transient increase in firing rate in response to each individual tone making up the stimulus sequence, with prominent adaptation by the directly preceding input. As expected, since the groups of tones in the stimulus itself do not match the periodic structure of the beat, the brainstem response shows a negative beat index (beat index = -0.15), indicating that a weakly-periodic input is not expected to elicit a prominent representation of the beat in the subcortical auditory nuclei.

A negative beat index is also obtained when the measure is computed directly from acoustic features of the rhythmic sequence known to strongly drive neural responses along the auditory pathway - from the auditory periphery to the primary auditory cortex - namely the acoustic envelope (beat index = −0.15) and spectral flux (beat index = −0.24). Overall, these negative beat index values confirm the dissimilarity between acoustic vs. beat structure that characterizes weakly-periodic rhythms.

### Beat representation is enhanced already in the primary auditory cortex

We then asked whether there was an enhancement of beat periodicity at the first cortical stage of sound processing, compared to the subcortical representation. To test this, we considered electrode contacts implanted along the Heschl’s gyrus (HG, or transverse temporal gyrus), typically considered as an anatomical landmark for the primary (or “core”) auditory cortex [38,47–49]. Indeed, 35 out of 36 implanted contacts showed a significant response to the stimulus, with a remarkable signal-to-noise ratio (SNR, quantified as responsiveness z-score, mean = 32.64, 95% CI [22.62, 42.66]). Next, we measured the prominence of beat periodicity in each of these responsive contacts. As shown in Fig 2, the beat index was significantly higher compared to the simulated brainstem response (one sample t-test, t_34_ = 5.32, P < 0.001).

**Fig 2.**
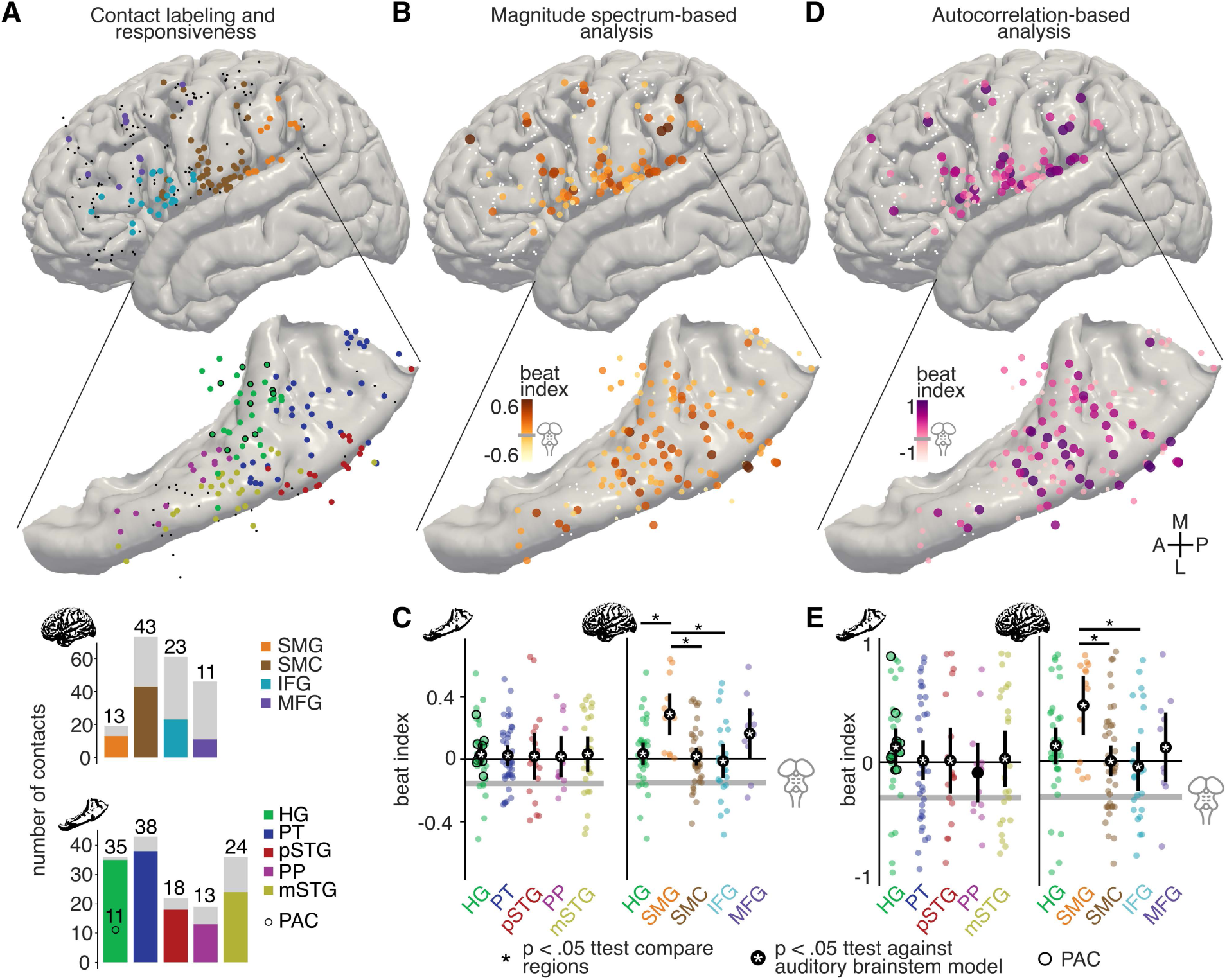
Anatomical parcellations and spatial distribution of beat index. (A) Anatomical labeling. Responsive recording contacts are shown in color. Locations of non-responsive contacts are indicated with black dots. For visualization, all contacts were projected to the cortical surface. Contacts in the right hemisphere were symmetrically projected across the midline. Pial surface with contacts localized in associative regions is shown on the top. The superior surface of the temporal lobe is shown below, with the corresponding auditory cortex contacts. Note that the nonlinear warping to a common template distorts the geometry of depth-electrode shafts (see S1 Fig for visualizations of electrode coverage on individual brains). The recording contacts labeled as primary auditory cortex based on a combination of anatomical and functional criteria are highlighted with a black outline. Bar plots visualize the total number of contacts (in gray) and the number of sound-responsive contacts (in color, also indicated above each bar) implanted in each region. (B) Spatial distribution of beat index values obtained from the magnitude spectrum of responses to the weakly-periodic rhythm. Non-responsive contacts are shown as white dots. The value obtained from the simulated brainstem response is indicated by a horizontal gray line in the color bar. The color scale is adjusted to highlight the relative differences between brain regions. (C) Prominence of beat periodicity across regions captured from magnitude spectrum. Only values from responsive contacts are shown. Data from auditory regions are shown on the left, and data from higher-order associative regions (along with HG included for comparison) are shown on the right. Each colored point represents a single recording contact. Means and 95% CIs are indicated by black points and error bars respectively. Regions where the observed beat index was significantly higher than in the simulated brainstem are highlighted with a white asterisk. Significant differences between regions (linear mixed-effects model, P < 0.05) are indicated by black asterisks. (D-E) Same as (B) and (C) but for beat index obtained from the autocorrelation function of each responsive contact.

**Fig 3.**
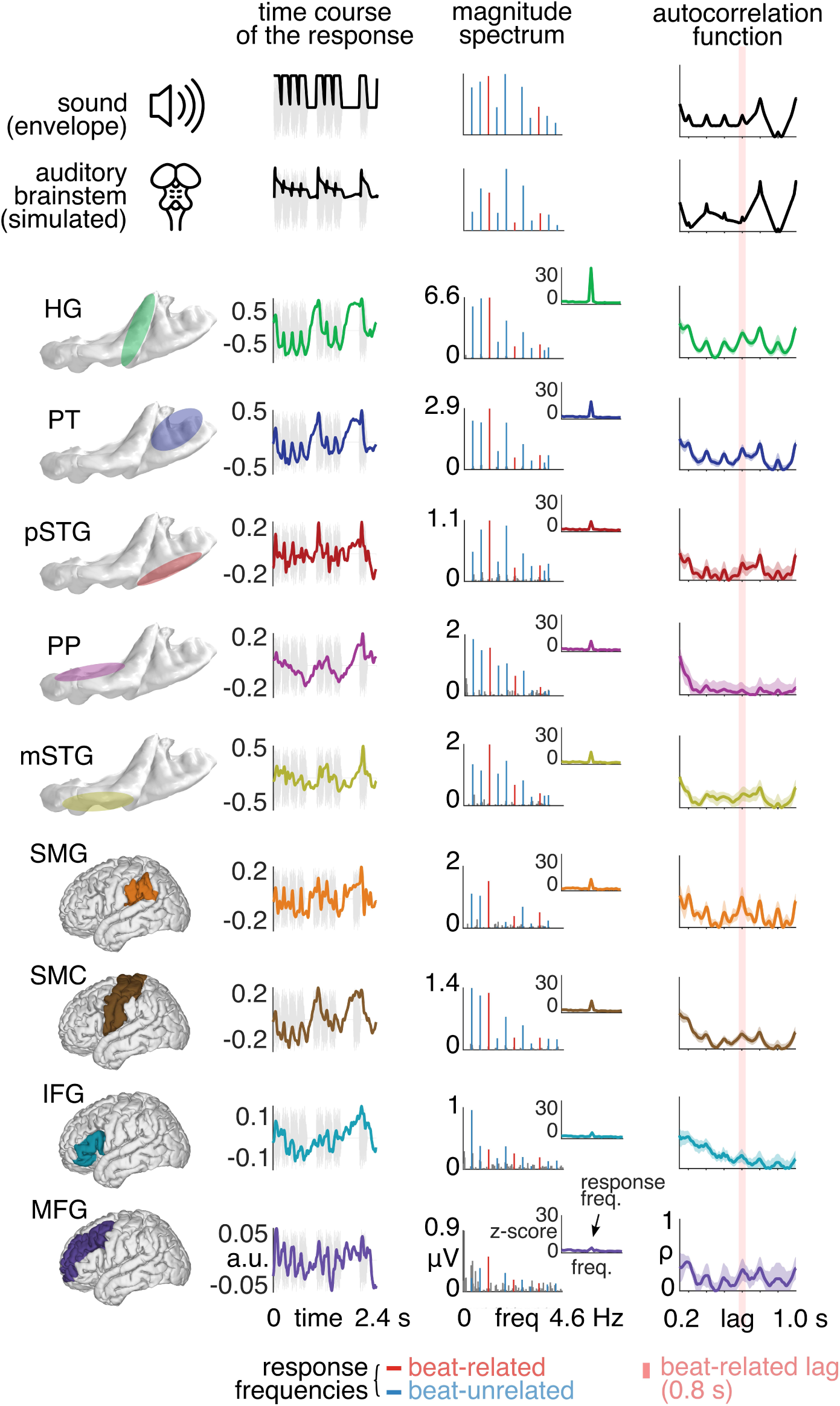
Summary of the response to the weakly-periodic rhythm in each anatomical region. The magnitude spectrum and autocorrelation function were averaged across all responsive contacts in the given region. The time domain response was estimated using Principal Component Analysis (PCA) to prevent sign cancellations across recording contacts. (Left) Average response time course to one repetition of the rhythmic pattern. (Middle) Magnitude spectrum. Frequencies related to the beat periodicity are shown in red and beat-unrelated frequencies used for magnitude normalization are shown in blue. (Right) Autocorrelation function. The lag corresponding to the consistently perceived beat period is indicated by a pink bar.

While HG constitutes a typical landmark to determine the approximate location of the primary auditory cortex, predicting functional properties from gross cortical anatomy poses a challenge, particularly in the supratemporal plane, which shows considerable inter-individual differences [47,50–52]. To further ensure the selected contacts were indeed capturing activity from the auditory core, we used a physiological criterion adopted in a number of previous studies. Specifically, we selected a subset of responsive contacts in the Heschl’s gyrus, that showed significant high-frequency locking above 130 Hz (see, e.g., [53–55] for a similar approach). This was done by evaluating whether the contact showed a significant frequency-following response at frequencies elicited in the cochlea by the partials making up the individual sounds in the stimulus sequence (see Methods). Out of the 35 responsive contacts, 11 showed significant high-frequency locking and were thus considered to capture activity from neurons in the primary auditory cortex (see Figs 2, and S1). Critically, even under such strict selection criteria, we still observed significant enhancement of beat periodicity compared to the simulated brainstem response (one sample t-test, t_10_ = 6.4, P < 0.001).

Together, these findings offer robust evidence that already at the first stage of cortical processing, the neural activity does not only reflect acoustic features but begins to gradually enhance beat-related perceptual information.

### Comparable beat representation across regions in the auditory cortex

Next, we asked whether the representation of beat periodicity may be further sharpened in secondary auditory areas, beyond the primary auditory cortex. To this end, we examined responses from contacts localized in several subfields of the temporal lobe auditory cortex, extending across the superior temporal plane and the adjacent lateral superior temporal gyrus (Fig 2A). We used anatomical divisions of the auditory cortex based on previous studies [38,56,57], which, besides HG, included the planum temporale (PT), planum polare (PP), posterior STG (pSTG) and middle STG (mSTG). As shown in Figs 2B and 2C, the beat index was comparable across auditory cortex regions (linear mixed-effects model, P = 0.48, BF_10_ = 0.13). This indicates that, rather than being further sharpened, the beat representation seems to be enhanced to a similar extent across the primary and secondary auditory cortex. Indeed, similar to the primary auditory cortex, the beat periodicity was significantly increased in each auditory region when compared to the simulated brainstem response (one sample t-tests, Ps < 0.05, FDR corrected, see S1 Table).

### Neural responses become strongly periodic in supramarginal gyrus, but not frontal associative regions

Even though the beat periodicity was significantly enhanced in the auditory cortices as opposed to the auditory brainstem model, in absolute terms, this periodicity did not prominently stand out in the temporal structure of the elicited responses. Indeed, none of the auditory cortices showed a significantly positive beat index (one sample t-tests, Ps > 0.05, FDR corrected, see S2 Table). Hence, it remains possible that for stimuli without periodic acoustic landmarks, the beat only fully emerges in higher-level associative regions that are interconnected with the temporal lobe auditory cortex. To test this, we considered a set of frontal and parietal regions that have been previously associated with rhythm processing and contained at least 10 sound-responsive contacts (Fig 2A). We used a linear mixed-effects model to compare the beat index across these regions. To directly compare associative regions with the auditory cortex, we also included data from HG in the model.

There were significant differences in beat index across regions (linear mixed-effects model, P = 0.002, BF_10_ = 11.28, Figs 2B and 2C). Post-hoc comparisons revealed increased beat index in the supramarginal gyrus (SMG) compared to the inferior frontal gyrus (IFG, P = 0.002) as well as the sensory-motor cortex (SMC, P = 0.006). In addition, SMG showed larger beat index compared to the HG (P = 0.02). In line with these results, while all tested regions showed a significantly enhanced beat periodicity compared to the biomimetic brainstem model (Ps < 0.05, FDR corrected, see S1 Table), SMG was the only region where the beat index was significantly greater than zero (P = 0.003, FDR corrected, see S2 Table). These results indicate a pivotal role of the SMG in the gradual transformation from acoustic features towards an abstract representation of a beat category.

One region that has been extensively reported as a core component of the beat processing network is the medial premotor cortex (MPC), comprising supplementary and presupplementary motor areas (SMA and preSMA, respectively) [21,22,26]. However, in our dataset, only 7 contacts (across 3 participants) showed a significant response to the sound input out of the 33 contacts implanted in the MPC across 4 participants. Even though this was below the criterion for inclusion of a region in our analysis, we still explored responses from the responsive MPC contacts to characterize their temporal dynamics. As shown in S2 Fig, the MPC responded to the stimulus sequence with slow smooth fluctuations, which is apparent in its magnitude spectrum, dominated by the first two harmonics of the rhythm repetition rate. These frequencies reflect a response that consistently repeats over the span of the rhythmic pattern, but not necessarily over the time span of the faster embedded beat period. Instead, the smooth response across MPC contacts appeared to track the groups of tones making up the rhythmic stimulus. Indeed, the beat index obtained from the magnitude spectrum was not significantly higher than the simulated brainstem response (one sample t-test, t_6_ = 0.84, P = 0.43).

### Strongly-periodic rhythm confounds representation of acoustic features and beat

Our findings so far reveal a gradient of transformation, whereby subcortical auditory nuclei are primarily driven by acoustic features, while beat periodicity is progressively enhanced along the auditory pathway, peaking in the inferior parietal cortex, which is bidirectionally connected with both auditory and frontal motor regions. Critically, these observations were made possible by our use of a weakly-periodic rhythmic stimulus, which allowed us to disentangle acoustic encoding from the perceptually relevant representation of beat periodicity.

To assess the potentially misleading influence of using stimuli that fail to control for this acoustic confound, we recorded intracerebral EEG responses while the same participants listened to a strongly-periodic rhythm. As with the weakly-periodic rhythm used above, this stimulus has been shown to reliably elicit beat perception with a period of 0.8 s in Western adults [8,9,13]. However, in this case, the temporal arrangement of the tones was closely aligned with the perceived beat, such that salient acoustic landmarks occurred at the beat period (S3A Fig).

As expected, already the simulated brainstem response was characterized by a highly prominent beat periodicity (beat index = 0.89), as was the acoustic envelope (beat index = 0.79) and spectral flux (beat index = 0.99) of the stimulus sequence. This prominent beat periodicity was maintained across temporal lobe auditory cortices (no difference between regions, linear mixed-effects model, P = 0.25, BF_10_ = 0.14), all showing a significantly positive beat index (one sample t-tests, Ps > 0.05, FDR corrected). In fact, while the beat index remained large, it was significantly smaller compared to the brainstem (one sample t-tests, Ps > 0.05, FDR corrected), likely due to the combination of a ceiling effect and noise (as compared to the subcortical response simulated with a biomimetic model rather than recorded in vivo) [7].

Considering areas beyond the auditory cortex, we observed significant differences between regions (linear mixed-effects model, P < 0.001, BF_10_ > 1000), driven by lower beat index in the IFG compared to HG (P < 0.001), SMC (P < 0.001), and SMG (P < 0.001). In addition, beat index was marginally smaller in the MFG compared to HG and SMG (P = 0.08 for both). This was expected based on the properties of the beat index variable, which has been shown to be pulled away from the true value towards zero as more and more noise is added to the response signal [7]. Indeed, we found a similar pattern of differences when comparing SNR across the same regions (linear mixed-effects model, P < 0.001, BF_10_ > 1000), with smaller SNR in the IFG and MFG as compared to SMG and SMC (Ps < 0.001) and an order of magnitude larger responsiveness in the HG compared to all higher-level associative regions (Ps < 0.001) (see also S5 Table and S3 Fig). Furthermore, pooling sound responsive contacts across all regions analyzed in the current study (i.e., all auditory and associative regions of interest), we observed a significant positive relationship between SNR and the beat index (linear mixed-effects model, β = 0.22, t_225_ = 5.33, P < 0.001, BF_10_ > 1000), confirming that contacts with overall smaller response compared to the noise floor tended to show smaller beat index.

Could regional differences in SNR explain the variation in beat index observed when participants listened to the weakly-periodic rhythm? If this were the case, one would expect a negative relationship between sound responsiveness and the beat index. Indeed, as the simulated auditory brainstem representation—unaffected by noise—exhibited a beat index below zero, decreasing SNR should bias the beat index toward zero, yielding a negative association between sound responsiveness and beat index across contacts [7]. However, these two variables were not significantly correlated (linear mixed-effects model, β = 0.05, t_223.78_ = 1.48, P = 0.14, BF_10_ = 0.16). Together, these results further support the interpretation that the selective enhancement of beat periodicity observed specifically for the weakly-periodic stimulus cannot be explained solely by regional differences in SNR.

### Excluding potential confounds based on waveform shape

In the analyses above, we relied on the magnitude spectrum as a highly sensitive measure to capture differences in the prominence of beat periodicity across signals, including both simulated firing-rate dynamics in the inferior colliculus and cortical activity measured with intracerebral EEG. However, the beat index derived from the magnitude spectrum has been shown to exhibit lower specificity, particularly when comparing signals that differ in the shape of their periodically recurring waveform [58]. To address this limitation, we conducted a complementary analysis using a novel autocorrelation-based approach that provides an estimate of beat periodicity that is invariant to the shape of the recurring signal [7]. This method quantifies the self-similarity of a signal after it is shifted by a lag corresponding to the beat period and compares this to the self-similarity obtained with other lags that are not related to the beat period. A standardized contrast between these two quantities is reflected in the resulting autocorrelation-based beat index.

First, we examined responses to the weakly-periodic stimulus. Autocorrelation-based analysis of the biomimetic brainstem model confirmed that beat periodicity was not prominent in the subcortical auditory representation (beat index = −0.32). Comparable lack of beat prominence was observed in the acoustic envelope (beat index = -0.27) and spectral flux (beat index = -0.35) of the stimulus sequence, corroborating the dissociation between acoustic features and the consistently perceived beat period, in line with the magnitude-spectrum analysis above.

Critically, compared to the simulated brainstem response, beat was significantly enhanced in the primary auditory cortex identified on the basis of anatomical landmarks (one-sample t test, t₃₄ = 5.62, P < 0.001), and this effect persisted when the contacts in the primary auditory cortex were further constrained using a stricter functional localization criterion (one-sample t test, t₁₀ = 5.73, P < 0.001). There was no further enhancement of the beat periodicity across non-primary auditory fields in the temporal lobe (linear mixed-effects model, P = 0.28, BF_10_ = 0.19, Figs 2D and 2E), and the beat index remained significantly above the brainstem model across all auditory regions except of PP (Ps < 0.05, FDR corrected, S3 Table). Comparisons among higher-level associative regions revealed significant differences (P = 0.02, BF_10_ = 4.02), with SMG showing higher beat index than IFG (P = 0.02) and SMC (P = 0.02). Although the SMG did not differ significantly from HG or MFG (P = 0.14 for both), it was the only region that showed a beat index significantly above zero (P = 0.01, FDR corrected, S4 Table), indicating strongly periodic neural responses to the weakly-periodic rhythm in this associative region. Finally, despite the small number of responsive MPC contacts, exploratory analyses showed that the beat index in this high-order motor planning area was numerically above the simulated brainstem representation (t_6_ = 2.34, P = 0.06) but clearly not significantly above zero (t_6_ = −1.42, P = 0.2).

The autocorrelation-based analysis further corroborated the inherent confound between acoustic features and beat periodicity for the strongly-periodic rhythm. Consistent with the magnitude spectrum analysis, the simulated brainstem response exhibited a saturated beat index (> 0.999), which remained significantly positive across all examined regions—including auditory and associative cortices (Ps < 0.05, FDR corrected)—with the exception of IFG and MFG. Consistent with the magnitude spectrum analysis, this was likely related to the differences in SNR (see S5 Table), which was positively correlated with the beat index across responsive contacts for the strongly-periodic rhythm (linear mixed-effects model, β = 0.43, t_223.6_ = 6.01, P < 0.001, BF_10_ > 1000). Critically, we also confirmed the significant positive relationship between SNR and beat index for the weakly-periodic rhythm (β = 0.21, t_223_ = 2.8, P = 0.006, BF_10_ = 0.38), indicating that rather than converging towards the negative beat index found in the lower-level subcortical representation, contacts with greater SNR tended to show more positive beat indices.

Taken together, the autocorrelation-based analysis corroborated the results based on the magnitude spectrum analysis, highlighting gradual enhancement of beat periodicity across cortical regions and the special role of SMG, revealed when the system is stimulated with a weakly-periodic rhythmic input.

### Local referencing scheme shows lower SNR and less sensitivity to capture progressive enhancement of beat periodicity beyond auditory cortex

Thus far, all results reported above were obtained using a common average reference. However, given that different referencing schemes may differentially emphasize or attenuate particular aspects of neural responses [59], we repeated the analyses for the weakly-periodic stimulus using a bipolar reference (see <u>S1 Text</u> for further details). Overall, we found that isolating highly local activity using bipolar referencing yielded a qualitatively similar pattern of results to the common average referencing, and successfully captured periodization between the subcortical and cortical stages of the auditory pathway. However, this montage appeared less sensitive to further transformations in the parietal cortex, likely related to the generally lower SNR after bipolar referencing (S4 Fig), which is known to attenuate spatially extended sources [59,60], thus particularly affecting high-amplitude low-frequency activity that has been shown fundamental to capturing neural transformation processes that underlie beat perception [8,14,20,61].

## Discussion

This study combines intracerebral EEG with biomimetic modeling to explore beat processing in the human brain. It provides direct evidence that a network of cortical regions responds to rhythm by gradually amplifying beat-related perceptual representations. Specifically, this transformation does not occur subcortically but emerges in the primary auditory cortex. In addition, this enhanced representation of the beat is also found in frontal associative regions, and is further sharpened in the inferior parietal cortex, yielding neural activity that is strongly periodic in response to a weakly-periodic rhythm. Together, these findings pinpoint the functional role of regions along the dorsal auditory pathway in supporting beat processing.

### Primary auditory cortex, but not auditory brainstem, shows enhanced beat representation beyond acoustic features

We show that, relative to a subcortical auditory representation, beat periodicity is significantly enhanced in the primary auditory cortex, corroborating previous intracerebral EEG work [20]. Although the present data cannot definitively rule out beat enhancement in the auditory brainstem emerging from mechanisms not captured by our biomimetic model, several independent observations make this interpretation unlikely. In particular, our findings are consistent with surface EEG measurements of brainstem responses in humans [9] and converge with electrophysiological data showing an absence of beat enhancement in the inferior colliculus of rodents when stimulated with rhythmic auditory sequences based on the same weakly-periodic pattern used in the current study [58; see their Figure S2].

Thus, in the broader context of previous work, our findings provide strong evidence that in humans (i) cortical processing is required to abstract from stimulus-driven acoustic encoding toward a more invariant representation of the beat, and (ii) this transformation emerges already at the earliest cortical stages of auditory processing. This interpretation is consistent with a growing body of surface EEG studies showing that beat-relevant transformation beyond acoustic features arises automatically during orthogonal tasks [8,9,13,14,19,62] and early in development (e.g., in 6-month-old infants) [17], while being at least partially attenuated during sleep [18].

### Inferior parietal cortex as a key region of the dorsal auditory pathway that supports abstract beat representation

Notably, we show that while the enhancement of beat periodicity in the auditory cortex is statistically significant when compared to the simulated auditory brainstem response, the magnitude of this transformation is arguably modest across the temporal lobe regions, and, similarly, across frontal associative regions. In contrast, the beat index becomes positive in the supramarginal gyrus (SMG, part of the inferior parietal cortex). This result reveals an invariant representation of the beat emerging in this parietal region in the form of strongly periodic neural activity.

Within influential predictive-processing accounts of beat perception [22,63], the parietal cortex is proposed to act as an interface between beat-based temporal predictions generated in frontal motor regions and the sensory representations encoded in auditory cortex. In this framework, parietal areas are thought to compute mismatches between predicted and incoming sensory signals and to relay these prediction errors back to motor regions to update the internal beat model [5,64]. While our study did not explicitly target prediction-error computations, the observed results suggest that SMG may contribute to beat perception by associating a perceptually relevant timing template with the incoming sensory input, beyond solely computing mismatches between predictions and sensory evidence [64]. In this respect, the inferior parietal cortex appears as a plausible candidate to support the transformation from largely acoustic encoding to higher-level perceptual representations, given its anatomical position at the intersection of auditory, somatosensory, and motor-related systems [65]. This higher-order associative cortical area would thus enable multimodal integration as well as plasticity and learning that have been shown to shape beat representations [66–70].

What could be the mechanisms underlying the emergence of such an abstract representation of the beat? One could speculate that this higher-level perceptual representation may emerge from the interaction between slower and faster periodic cues present in the sensory input. Specifically, for the weakly-periodic stimulus used here, these periodicities corresponded to the repetition of the rhythmic pattern (2.4-s period) and the recurrence of the smallest inter-onset interval (0.2-s period) within groups of tones in the stimulus sequence. These slower and faster acoustic periodicities could interact through a process of multiscale temporal scaffolding, eliciting representation of an intermediate pulse layer corresponding to the beat targeted in the present study and embedded into this holistic nested configuration (i.e. musical meter) [14,19].

An influential neurocomputational framework that offers a mechanistic account of such multiscale temporal scaffolding is the neural resonance theory [24,25]. This theory posits that, in response to weakly-periodic inputs, beat periodicity can be selectively amplified through nonlinear coupling between two bidirectionally connected oscillatory networks: one network models auditory cortical dynamics while the second one is hypothesized to reflect motor cortical dynamics. Notably, simulations of this model indicate that beat enhancement arises primarily within the motor network [11], especially when beat-related periodicity is weak or absent in the sensory input. Our intracerebral data thus provide nuances to this view by suggesting that the putative second oscillatory network proposed in this theory may involve the parietal cortex, at least in the absence of task demands. This interpretation is broadly consistent with recent magnetoencephalography work that used source reconstruction to locate strongest beat-related periodization in the superior temporal cortex rather than in frontal motor areas [12]. In addition, the parietal cortex is well positioned to support such a function, as it is broadly implicated in sensory–motor transformations underlying auditory–motor integration for motor control [71–73].

### Similar beat enhancement across temporal lobe auditory cortices

The temporal lobe auditory cortex is often conceived as hierarchically organized, with increasingly abstract representations emerging from primary to non-primary, higher-order areas [49,71,74,75]. Within this anatomical–functional framework, neural activity in the primary auditory cortex is expected to closely track stimulus features, with progressive transformations occurring in surrounding, non-primary fields [40,76–78]. Contrary to this prediction, our data did not reveal a clear hierarchical gradient along the superior temporal lobe. Instead, beat representation was comparably prominent across auditory regions relative to the simulated subcortical response. This pattern is consistent with recent intracranial EEG studies reporting broadly distributed representations of higher-level attributes of speech [38,39,79,80] and musical pitch sequences [37,41] throughout the superior temporal lobe.

Notably, our findings of broadly distributed representations of the beat across temporal lobe auditory cortices can still be explained by a hierarchical account, whereby a critical transformation emerging locally in the primary auditory cortex is subsequently inherited by downstream secondary regions. Alternatively, the selective enhancement of beat periodicity may arise within a broader recurrent network spanning the entire temporal lobe. This alternative interpretation aligns with ongoing debates about the validity of strictly hierarchical models [42,43], and with accumulating evidence for parallel and distributed processing within the supratemporal plane and adjacent superior temporal gyrus [38,81]. Future studies using for example direct electrical stimulation will be necessary to further elucidate the directionality of functional connections underlying the spatially widespread beat-related periodization observed across the auditory cortex [82].

### Beat enhancement is comparable across auditory and frontal lobe cortices

Beyond auditory cortices, our results show that the beat representation was also significantly and comparably enhanced in higher-order associative regions of the frontal lobe. The presence of sound-responsive contacts throughout the frontal associative cortex has been reported in prior intracerebral EEG studies showing widespread sound-evoked frontal activity, including responses to musical rhythm [83] and speech [42,84–86]. This observation also aligns with influential neuroimaging studies of rhythm processing, which have reported broad BOLD activations elicited by rhythmic sounds in premotor cortex, spanning both lateral and medial regions [26,27,87–89]. Critically, the present study advances beyond prior work by not only testing whether a given site reliably *responds* to rhythmic input relative to baseline, but by directly assessing the *beat periodicity encoded in that response*, leveraging a robust and validated methodological approach [6,7,34,35]. Our findings thus provide direct insight into the putative nature of the information carried by frontal motor activations, and the functional role of these areas in supporting beat perception.

These insights are particularly timely in light of growing evidence for rapid, direct projections from human auditory cortex to frontal associative regions [42], and the emerging view that the targeted frontal areas primarily encode relatively low-level acoustic features rather than highly abstracted “motor” representations [42,84,90,91]. Our findings are broadly consistent with this perspective, in that beat periodicity in frontal associative regions did not exceed that observed in temporal lobe auditory cortex.

Notably, these results appear difficult to reconcile with influential evidence for a central role of frontal motor areas in beat processing derived from electrophysiological recordings in non-human primates. This research has shown that periodic population dynamics in macaque premotor cortex encode the beat during synchronization–continuation tapping tasks [29,30] and covert metronome tracking [31]. In light of this body of work, our results provide a unique opportunity to probe whether beat-related periodic dynamics also emerge in the human motor cortex during attentive listening, without an explicit motor task and extensive training, as predicted by prominent theoretical models of beat processing (for a review, see [5,23]). Interestingly, while our data showed a significantly enhanced representation of the beat in sound-responsive frontal contacts, these regions did not show strongly periodic neural activity abstracted out from acoustic encoding, as observed in the parietal cortex. Based on this result, we speculate that rather than being necessary to support beat representations and perform beat categorization functions, frontal motor regions may play a modulatory role [92], and become engaged depending on task demands, particularly when the beat representation is used explicitly for movement coordination [87,93,94], temporal predictions [31], and perceptual judgements [26,95]. Such a model closely parallels contemporary views on the role of the motor system in speech perception [72,91,96]. At the same time, it should be acknowledged that our frontal lobe coverage was relatively sparse. Studies with larger cohorts and more targeted electrode coverage will be required to more precisely delineate the functional contribution of frontal associative regions to beat processing. The present study nonetheless provides a principled framework for addressing these questions in future work.

### Conclusions

Overall, our findings contribute to a growing body of evidence suggesting that the neural bases of beat categorization may involve multiple, potentially interacting mechanisms implemented across partially distinct functional networks. These include neural adaptation [58], frequency tuning [8,9,20], nonlinear mechanisms [6,12,19], as well as associative learning mediated by neural plasticity [66,68,69,97,98] and volitional control [15,92,99,100]. Together, these processes may progressively transform neural representations from sensory-driven responses toward invariant yet flexible beat templates shared across individuals to support human music practices [101].

## Methods

### Participants

The study included 13 patients (5 females, mean age: 32 ± 12 years, 8 right-handed) undergoing clinical intracerebral evaluation with depth-electrodes (StereoElectroEncephaloGraphy, SEEG) for refractory partial epilepsy at the Epilepsy Unit of the University Hospital of Nancy (Nancy, France; 12 patients) and the Department of Neurology of Saint Luc University Hospital (Brussels, Belgium; 1 patient). The implantation of intracerebral electrodes was performed solely for clinical purposes, as part of presurgical evaluation to identify seizure foci. All participants reported no history of hearing or movement disorders. Participants were included in the study if they had at least one intracerebral electrode implanted in the auditory cortex or in the medial frontal cortex. All participants provided written consent to take part in the study, which was conducted according to the principles expressed in the Declaration of Helsinki and approved by a national ethics committee certified by the French Ministry of Health (Institutional Review Board: IORG0009855) and Commission Ethique de l’Université catholique de Louvain (B403201316436).

### Stimuli

The stimuli consisted of a 2.4-s long rhythmic pattern that was seamlessly repeated 17 times to form a continuous 40.8-s long sequence. The pattern consisted of 8 identical sound events arranged on an isochronous grid of 12 time points separated by 0.2 s. This arrangement can be represented as [xxxx.xxx..x.] where “x” stands for a grid position with a sound event and “.” marks an empty grid position. This particular rhythm was selected for two reasons. First, numerous studies have shown that this rhythm induces perception of a periodic beat, at a rate generally converging across Western participants toward 0.8 s (i.e., four grid intervals, 4 × 0.2 s) [8,9,13,14,61]. Second, the temporal structure of this rhythm does not clearly align with any plausible periodic beat. That is, the acoustic features of the stimulus sequence based on this “weakly-periodic” rhythm do not exhibit prominent periodicity at the rate of the typically perceived beat. To demonstrate the issue of confounding acoustic features and the perceived beat, we additionally created sequences based on a “strongly-periodic” rhythmic pattern, where the 8 identical sound events were rearranged to clearly match the periodic beat at the rate of 0.8-s ([xxx.xxx.xx..]).

The sound event corresponded to a 200-ms long complex tone made of 3 pure tone partials, with a 10-ms onset and 50-ms offset linear ramp. The partial frequencies corresponded to 234.1, 311.3 and 490.2 Hz for 10 participants, 243, 398, and 566 Hz for 3 participants. One participant additionally listened to sequences where the sound event corresponded to white noise with the same duration and ramp parameters as the tone event.

### Procedure

The stimuli were presented binaurally via insert earphones (ER4-SR, Etymotic Research, Elk Grove Village, IL) connected to a Fireface UC audio interface (RME Audio, Haimhausen, Germany) at a comfortable hearing level (around 70 dB). The experiment was implemented using PsychToolbox, version 3.0.14 [102]. Participants performed the experiment while comfortably seated in a hospital bed with their head resting on a support. They were instructed to relax, avoid any unnecessary head or body movement, and keep their eyes fixated on a single point in front of them to avoid any eye movements.

Participants were presented with stimulus sequences made of either the weakly-periodic or the strongly-periodic rhythm in alternation (order randomized across participants). They were instructed to avoid any movement and carefully listen to the sound. To further encourage attention to the stimuli, participants were asked to report any subtle changes in the tempo of the rhythm at the end of each trial. No actual tempo changes were present in any of the stimuli. The onset of each sequence was triggered by the experimenter when the participant felt ready for the following trial.

Each listening trial was followed by a tapping trial, whereby the same stimulus sequence was presented again, and participants were asked to tap their hand in time with the rhythm on a custom response box. For two participants the experiment only consisted of listening trials. The data from tapping trials are not analyzed in the current study.

Among the 13 participants, participants heard the weakly-periodic sequence either 3 times (1 participant), 4 times (11 participants), or 8 times (1 participant). Similarly, participants heard the strongly-periodic sequence either 3 times (1 participant), 4 times (11 participants), or 7 times (1 participant). No participant had seizures in the 2 hours preceding the recordings. Stimulus polarity was inverted in half of the trials to minimize the electromagnetic artefact generated by the sound delivery system.

### Intracerebral electrode implantation and EEG recording

Intracerebral electrodes (Dixi Medical, Besançon, France) were stereotactically implanted within the participants’ brains for clinical purposes, to delineate their seizure onset zones and to functionally map the surrounding cortex for potential epilepsy surgery. Each intracerebral electrode consisted of a cylinder of 0.8 mm diameter and contained 5–18 independent recording contacts of 2 mm in length separated by 1.5 mm from edge to edge and by 3.5 mm center-to-center (for details about the electrode implantation procedure, see [103]). Intracerebral EEG was recorded with a 256-channel amplifier referenced to a midline prefrontal scalp electrode (FPz). The sampling rate was either 2048 Hz (11 participants), 1024 Hz (1 participant), or 512 (1 participant). Channels containing excessive artifacts were identified by visual inspection and excluded from further analysis. The data were re-referenced offline to the common average comprised of all electrodes. For a control analysis, bipolar reference was obtained for each recording contact by subtracting the signal from the directly adjacent contact located more laterally on the same intracerebral EEG electrode array. The anatomical location of the resulting bipolar channel was calculated as the midpoint between the active and reference contact.

### Contact localization

The exact anatomical location of each recording contact was determined by coregistration of post-operative non-stereotactic CT-scan with a pre-operative T1-weighted MRI. Across the 13 participants, a total of 110 electrode arrays were implanted in the auditory cortex, and the parietal and frontal associative regions of interest. These electrodes contained 452 individual recording contacts (i.e., in the gray/white matter; 266 contacts in the left hemisphere, 186 in the right hemisphere).

Three-dimensional reconstructions of the pial surface were created using an individual subject’s pre-operative T1 MRI scan in Freesurfer and anatomical regions of interest for each electrode were automatically assigned using the Desikan-Killiany atlas [104]. To account for the sparse coverage of intracerebral EEG, we created larger regions of interest corresponding to associative brain areas implicated in rhythm and timing by merging contacts across the Desikan-Killiany atlas labels. Specifically, “parsopercularis”, “parstriangularis”, and “parsorbitalis” were pooled to form the inferior frontal gyrus (IFG), “caudalmiddlefrontal” and “rostralmiddlefrontal” labels were merged into middle frontal gyrus (MFG), and sensory-motor cortex (SMC) was created by pooling contacts in the “precentral” and “postcentral” gyrus.

In addition to the automatic labeling, contacts in the supratemporal plane (STP) were labeled manually based on individual anatomical landmarks, following Hamilton et al. [38]. Likewise, contacts in the supplementary and presupplementary motor areas (SMA and preSMA, respectively) were localized in the medial frontal cortex superior to the cingulate sulcus. Specifically, contacts posterior to the vertical commissure anterior (VCA) line all the way to the vertical commissure posterior (VCP) line were labelled as SMA [105], and contacts anterior to the VCA line up to the genu of corpus callosum line were labelled as preSMA [106]. SMA and preSMA were together considered the medial premotor cortex (MPC). The spatial location of electrode arrays that had at least one recording contact in one of the regions of interest is shown in S1 Fig.

For the control analysis using bipolar reference, anatomical labels were determined based on the anatomical label of the active and reference contact. In cases where one contact was located outside the gray matter, the label corresponding to the gray-matter contact was used. If the active and reference contact were assigned to two different regions, the label for the active contact was used.

We warped electrode coordinates onto the cvs_avg35_inMNI152 template [107] using a combination of volumetric and surface registration [108]. For visualization purposes, each electrode was projected onto the reconstructed pial surface (see Figs 2A, 2B, and 2D).

### Intracerebral EEG signal processing and analyses

Intracerebral EEG signal processing was similar to previous studies using periodic stimulation [20,109,110]. First, a notch filter (filter width = 2 Hz, FFT filter) was applied to the continuous data to minimize contamination with line noise at 50 Hz (and harmonics up to 200 Hz).

Data were then segmented into epochs between 0 and 40.8 seconds relative to sound onset, thus encompassing the total duration of stimulation in each trial (i.e., exactly 17 repetitions of the 2.4-s long rhythmic pattern). The epochs were subsequently averaged in the time domain, separately for each rhythm, electrode, and participant. The time-averaged data were transformed into the frequency domain using fast Fourier transform (FFT), yielding a spectrum with resolution of approximately 0.02 Hz.

#### Quantifying responsiveness to sound

To evaluate whether an electrode contact showed a significant response to the rhythmic stimulation that stood out from the background noise level, we capitalized on the fact that the spectrum of any signal that is systematically repeated with a fixed repetition rate (i.e., periodically) will only contain peaks at specific frequencies corresponding to the repetition rate (i.e., f = 1/repetition period) and its integer multiples (i.e., harmonics, 2f, 3f, 4f, etc.) [7,111]. In the current study, the stimulus sequences were composed of a seamlessly repeated rhythmic pattern presented with an exact fixed repetition period (2.4 s). Therefore, the spectrum of any response systematically elicited across repetitions of the rhythm pattern is expected to contain responses at frequencies corresponding to f = [1 / pattern duration] and harmonics (i.e., f = 1/2.4 s = 0.417 Hz, 2f = 0.83 Hz, …). We considered the first 11 harmonics of the rhythm repetition rate as response frequencies in further analyses, based on our recent work showing that auditory EEG responses to the rhythmic sequences used here are concentrated in the low-frequency bandwidth below 5 Hz [8,14,20,61].

Following a procedure adopted in previous studies using periodic stimulation [112–115], we assessed the responsiveness of each electrode contact as follows: (i) the magnitude spectrum was cut into segments centered on each response frequency including 13 neighboring frequency bins (corresponding to 0.32 Hz) on each side; (ii) the magnitude values were summed across segments separately for each frequency bin; (iii) the summed magnitude spectrum was transformed into a z-score. The z-score was computed as the difference between the magnitude at the response frequency bin and the mean magnitude of surrounding bins (excluding two closest bins on each side of the response frequency bin), divided by the standard deviation of the magnitudes across the surrounding bins. A contact was considered responsive to sound if the z-score value (hereafter referred to as “responsiveness z-score”) computed for its response to the weakly-periodic stimulus sequence exceeded 2.3 (i.e., p < 0.01 one-tailed: signal > noise). The responsiveness z-score was used in further analyses as a continuous index of signal-to-noise ratio (SNR).

#### Identifying primary auditory cortex based on physiological criteria

To carry out an even more strict selection of contacts within the primary auditory cortex, we applied a functional localizer inspired by previous intracerebral EEG studies [53,55]. This localizer was based on prior evidence that neuronal populations in the heavily myelinated tonotopic “core” auditory cortex can phase-lock their activity to high-frequency inputs above 100 Hz, whereas this is not the case for responses from the surrounding non-primary fields [47,116,117]. Thus, electrodes capturing responses from the primary auditory cortex would be expected to show a significant frequency-following response at the partial frequencies composing the tones conveying the rhythmic inputs, as well as high-frequency (above 100 Hz) distortion products arising from interactions between these partials at the cochlear level [118]. Here, we only considered quadratic and cubic distortion products rather than partials, to ensure no contamination of the response with any remaining electromagnetic artefact of the sound delivery system. These distortion products corresponded to 256.1, 178.9, 156.9, and 132.4 Hz for 10 participants, and 155, 323, 168, and 230 Hz for 3 participants. Since the primary auditory cortex in humans is thought to be roughly located on the Heschl’s gyrus, we only analyzed activity of sound-responsive contacts delineated by this anatomical landmark. We quantified frequency-following of the distortion products by applying the same method used to identify sound-responsive contacts across the whole brain. The magnitude spectrum was cut into segments centered on each distortion product frequency. These snippets were summed and the magnitude at the distortion product bin was converted into a z-score using the surrounding bins. A contact was considered to capture primary auditory cortex activity if its distortion product z-score exceeded 2.3 (i.e., p < 0.01 one-tailed: signal > noise).

#### Quantifying prominence of beat periodicity using magnitude spectrum

To quantify the periodicity of the response at the rate of the beat, we first extracted baseline-corrected magnitudes at response frequencies. Baseline correction was done by subtracting the average magnitude at 13 surrounding bins on each side of the given response frequency (excluding two closest bins on each side to account for any remaining spectral leakage). Next, we selected a subset of response frequencies corresponding to the rate of the beat typically induced by the stimulus sequences in Western adults, and its harmonics (i.e. f_beat_ = 1/0.8 s = 1.25 Hz, 2f_beat_ = 2.5 Hz, and 3f_beat_ = 3.75 Hz). The relative prominence of these beat-related frequencies within the whole set of response frequencies quantifies the beat periodicity reflected in the response. To capture this prominence, we first standardized magnitudes across all 11 response frequencies using z-scoring (i.e., by taking the difference between the magnitude at each response frequency and the mean magnitude across all response frequencies, divided by the standard deviation of magnitudes across all response frequencies). Then, we averaged z-scores across beat-related frequencies to obtain an index of beat periodicity reflected in the response. Finally, the mean z-score at beat-related frequencies was scaled to obtain the beat index using the following equation

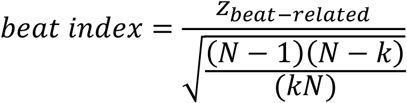

where N is the number of response frequencies (here N=11) and k is the number of beat-related frequencies (here k=3). This scaling ensured beat z-scores can only take values between -1 (indicating no beat periodicity, magnitude solely concentrated at beat-*unrelated* frequencies) and 1 (indicating perfect beat periodicity, magnitude solely concentrated at beat-*related* frequencies).

#### Quantifying prominence of beat periodicity using autocorrelation

To quantify the prominence of beat periodicity using autocorrelation, the time-averaged intracerebral EEG data were low-passed filtered at 30 Hz (2^nd^ order Butterworth filter) to isolate the low-frequency bandwidth of electrophysiological responses to acoustic rhythms [14,20]. To minimize contamination of the autocorrelation estimate with noise, we used a two-step noise-correction method as described in Lenc et al. [7]. First, we estimated the spectrum of the 1/f-like noise baseline using irregular resampling with scaling factors between 1.1 and 2 in steps of 0.05 (as implemented in the IRASA package; see [119]). The data were subsequently transformed into the frequency domain using FFT and the 1/f-like noise estimate was subtracted from the resulting complex spectrum. To further reduce the contribution of noise, we set to zero all complex Fourier coefficients at frequency bins that did not correspond to the harmonics of the rhythm repetition rate. Finally, we used the convolution theorem to obtain circular autocorrelation from the noise-corrected spectrum [120]. The resulting autocorrelation function reflects the similarity of the signal with a lagged version of itself, across a range of lags from 0 s to half of its duration, here corresponding to half the total duration of the rhythmic sequence (i.e. 40.8 s / 2 = 20.4 s). From this autocorrelation function, we extracted values at lags of interest corresponding to (i) beat periodicities (beat-related lags, 0.8 s and multiples up to the highest obtained lag) and (ii) control lags corresponding to periodicities where the beat was not perceived despite being compatible with the temporal arrangement of the sounds making up the rhythmic stimulus (beat-unrelated lags, 0.6, 1.0, 1.4, and their multiples). Overlapping beat-related and beat-unrelated lags were excluded. In addition, any multiples of 0.4 s were excluded from the set of beat-unrelated lags, since this periodicity (a duple subdivision of the beat) has been identified as part of the meter typically induced by the rhythms used in the current study [9,13]. After normalizing the autocorrelation coefficients across the whole set of lags using z-scoring, the periodicity of the response at the rate of the beat was quantified by averaging the coefficients across beat-related lags. The resulting beat-related z-score was scaled to fall between -1 and 1, using the same approach as for the magnitude spectrum analysis above.

#### Time-domain analysis

To visualize the time-domain dynamics of the response from each region of interest, the continuous data from each electrode was low-pass filtered at 20 Hz (4^th^ order Butterworth filter), segmented from the onset to the end of the stimulus sequence (0 to 40.8 s), and averaged across trials, separately for each electrode, rhythm, and participant. The data were then segmented into successive chunks corresponding to the rhythmic pattern duration (i.e., 2.4 s), averaged across the 17 resulting epochs, and down sampled to 128 Hz. To obtain the characteristic response time course from given region of interest, we took the first principal component after applying principal component analysis (PCA) to data from all electrodes in that region (i.e., collapsed across participants). PCA was used instead of average to prevent potential cancellation of responses across electrodes within a region due to polarity inversion.

### Biomimetic model of subcortical auditory responses

To estimate the neural representation of the rhythmic stimulus in the auditory brainstem, we simulated responses to the stimulus sequence using a biologically plausible model of subcortical auditory processing as implemented in the UR_EAR toolbox (version 2020b). First, a model of the auditory nerve was used to simulate responses from 128 cochlear channels with characteristic frequencies logarithmically spaced between 130 and 8000 Hz [46]. The parameters used for cochlear tuning matched data available from human subjects [121]. For each channel, 51 auditory nerve fibers were simulated with biologically plausible distribution of high, mid, and low-spontaneous-rate fibers [122]. The model provides faithful simulation of physiological processes associated with cochlear nonlinearities, inner hair cell transduction process, the synapse between the hair cell and the auditory nerve, and the associated firing rate adaptation. The simulated auditory nerve firing rates were fed into the same-frequency inhibition and excitation model (SFIE) used to simulate enhanced onset synchrony and the decreased upper limit for phase-locking to stimulus envelope in the ventral cochlear nucleus [44]. The default parameters in the UR_EAR toolbox were used, which were based on Carney et al. [45]. A second SFIE model was then used to simulate bandpass modulation filtering and enhanced onset responses of neurons in the inferior colliculus. The parameters were set to simulate inferior colliculus units with the best modulation frequencies separately at 2, 4, 8, 16, 32, and 64 Hz. The firing rates were summed across all channels and modulation frequencies. The obtained time course was taken as an approximation of the instantaneous firing rate over the course of the 40.8-s long stimulus sequence from a population of neurons in the inferior colliculus. This time course was then analyzed in the same way as the intracerebral EEG responses, using both the magnitude spectrum and autocorrelation-based approach, to quantify periodicity of the brainstem response at the rate of the beat.

In addition, we applied the same analyses to acoustic features of the stimulus sequence (envelope and spectral flux) that have been shown to strongly drive neural responses along the auditory pathway [123,124]. The broadband amplitude envelope was estimated using Hilbert transform. Spectral flux (i.e., changes in spectral content over time) was estimated by computing logarithmically compressed spectrograms in 50-ms windows (10-ms hop size), calculating the temporal derivative across successive windows, and setting negative values to zero to isolate increases in intensity [125].

### Statistical analyses

Contacts within the same region were grouped and collapsed across hemispheres. Hemispheric comparisons were not conducted due to the limited sampling of the right hemisphere, with fewer than 10 contacts available in key regions (including HG, PP, pSTG, and SMG). Beat index values from each region were compared to the corresponding value obtained from the biomimetic brainstem model or to zero using one sample t-tests. P-values were corrected for multiple comparisons using Benjamini-Hochberg false discovery rate correction (FDR) [126].

To statistically compare beat index values across brain regions we used general linear mixed-effects models implemented in the lme4 package [127] in R v4.5.2. Models were fitted using restricted maximum likelihood (REML) and compared using Type II Wald F tests (as implemented in the *car* package [128]). P-values were obtained using Kenward-Roger’s approximation for degrees of freedom. Models included a random intercept per participant to account for both inter-participants variability and non-independence of contacts within a participant. Posthoc multiple comparisons were computed using the *emmeans* package [129], and corrected for multiple comparisons using FDR.

Additionally, we assessed the relationship between overall sound responsiveness (SNR) and beat periodicity (derived from either the magnitude spectrum or autocorrelation) using linear mixed-effects models fit to all responsive electrode contacts pooled across regions. Since responsiveness z-score values were not normally distributed, we applied a log transformation. Beat index was modeled as the dependent variable, log-transformed sound-responsiveness z-score as a continuous fixed predictor, and participant as a random intercept. Furthermore, SNR was directly compared across brain regions, using a linear mixed-effects model with log-transformed sound-responsiveness z-score as a dependent variable, brain region as categorical fixed predictor, and participant as a random intercept. Finally, the same model was used to compare SNR between the common average and bipolar reference montage, using reference type (common average reference, bipolar) as an additional categorical fixed predictor. All electrode contacts were included when assessing differences across reference montages to ensure the validity of the comparison.

Bayes factors (BF₁₀) were additionally computed to quantify evidence for the alternative relative to the null hypothesis, using the BayesFactor package in R [130].

### Code accessibility

Numerical data and analysis code to reproduce the results and figures of this manuscript will be shared upon publication.

## Acknowledgements/Funding

This work was supported by the European Research Council (801872, 101228872) and the Marie Sklodowska-Curie programme (101148958). We are grateful to Giulia Liberati for her help with data collection at the Saint Luc University Hospital (Brussels, Belgium).

## Supplementary Materials

**S1 Table.** One-sample t-tests, comparing beat index obtained from the magnitude spectrum of the intracerebral EEG responses to the weakly-periodic rhythm with the value obtained from the simulated brainstem response.

| ROI | t | df | P |  |
| --- | --- | --- | --- | --- |
| HG | 5.32 | 34 | <0.0001 | *** |
| PP | 2.84 | 12 | 0.017 | * |
| PT | 5.42 | 37 | <0.0001 | *** |
| pSTG | 2.46 | 17 | 0.025 | * |
| mSTG | 3.42 | 23 | 0.004 | ** |
| SMG | 7.12 | 12 | <0.0001 | *** |
| SMC | 6.08 | 42 | <0.0001 | *** |
| IFG | 2.68 | 22 | 0.017 | * |
| MFG | 4.45 | 10 | 0.002 | ** |

\* $P < 0.05$ , \*\* $P < 0.01$ , \*\*\* $P < 0.001$ , FDR corrected

**S2 Table.**
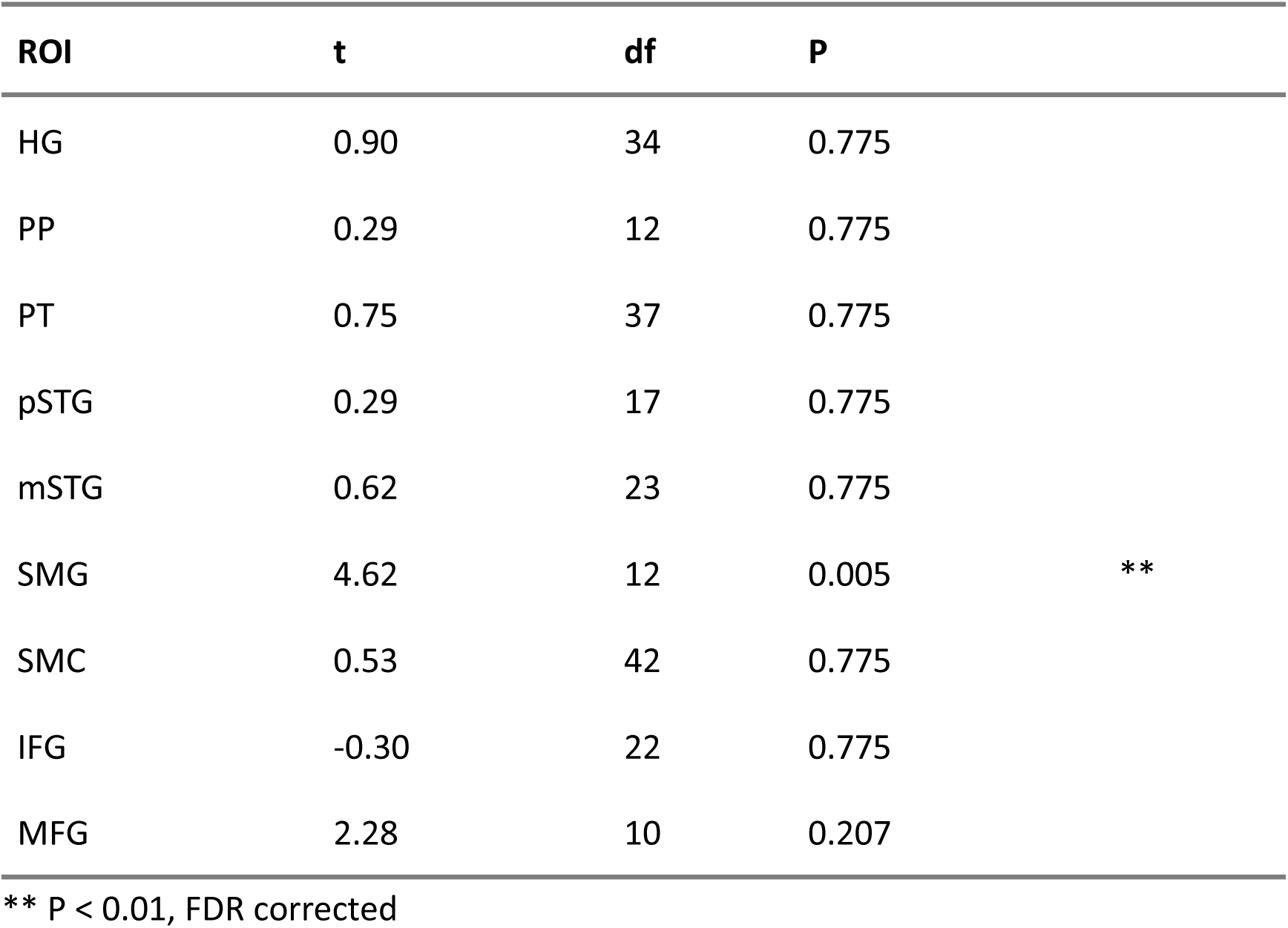
One-sample t-tests, comparing beat index obtained from the magnitude spectrum of intracerebral EEG responses to the weakly-periodic rhythm to zero.

**S3 Table.**
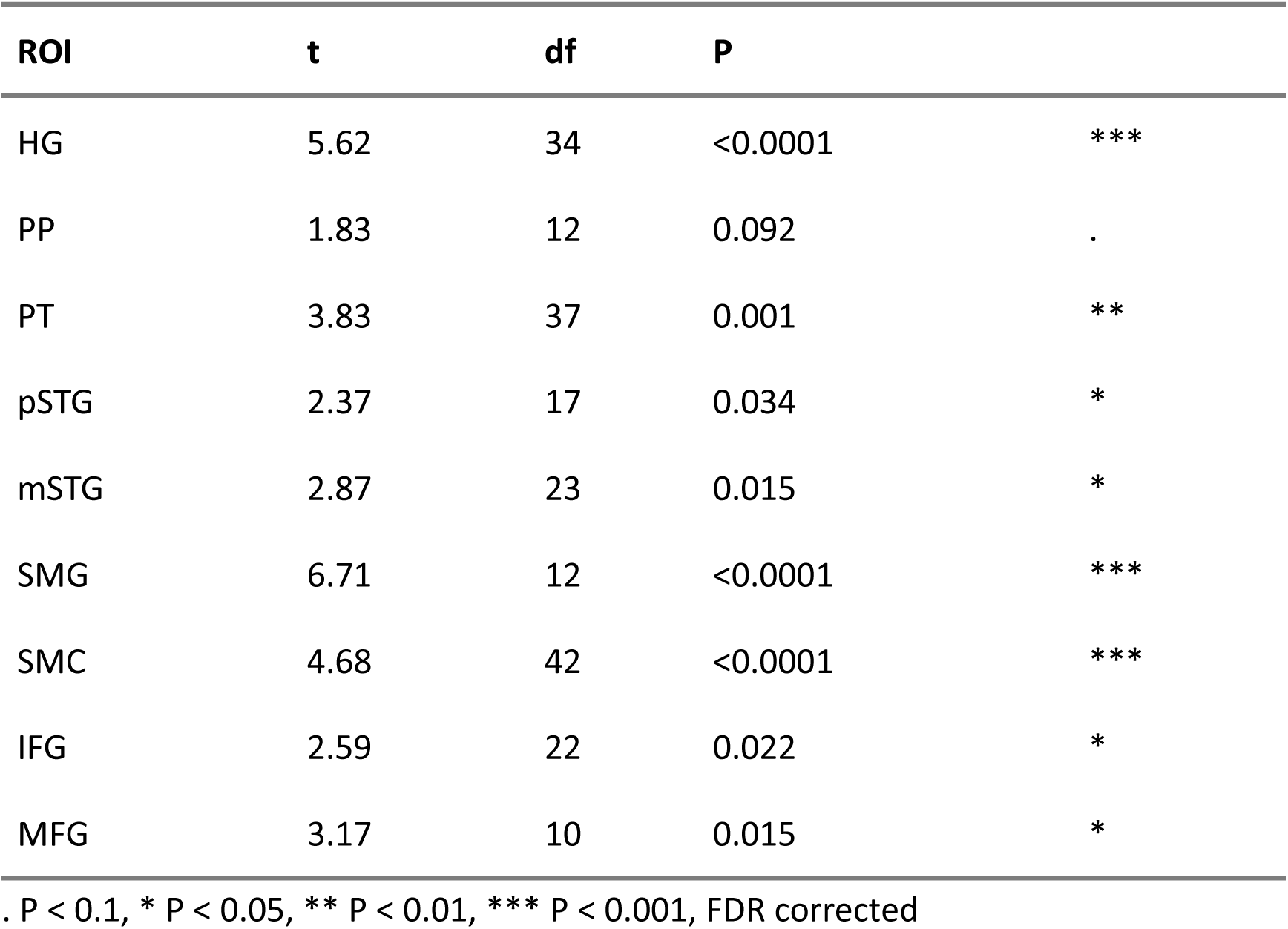
One-sample t-tests, comparing beat index obtained from the autocorrelation of intracerebral EEG responses to the weakly-periodic rhythm with the value obtained from the simulated brainstem response.

**S4 Table.**
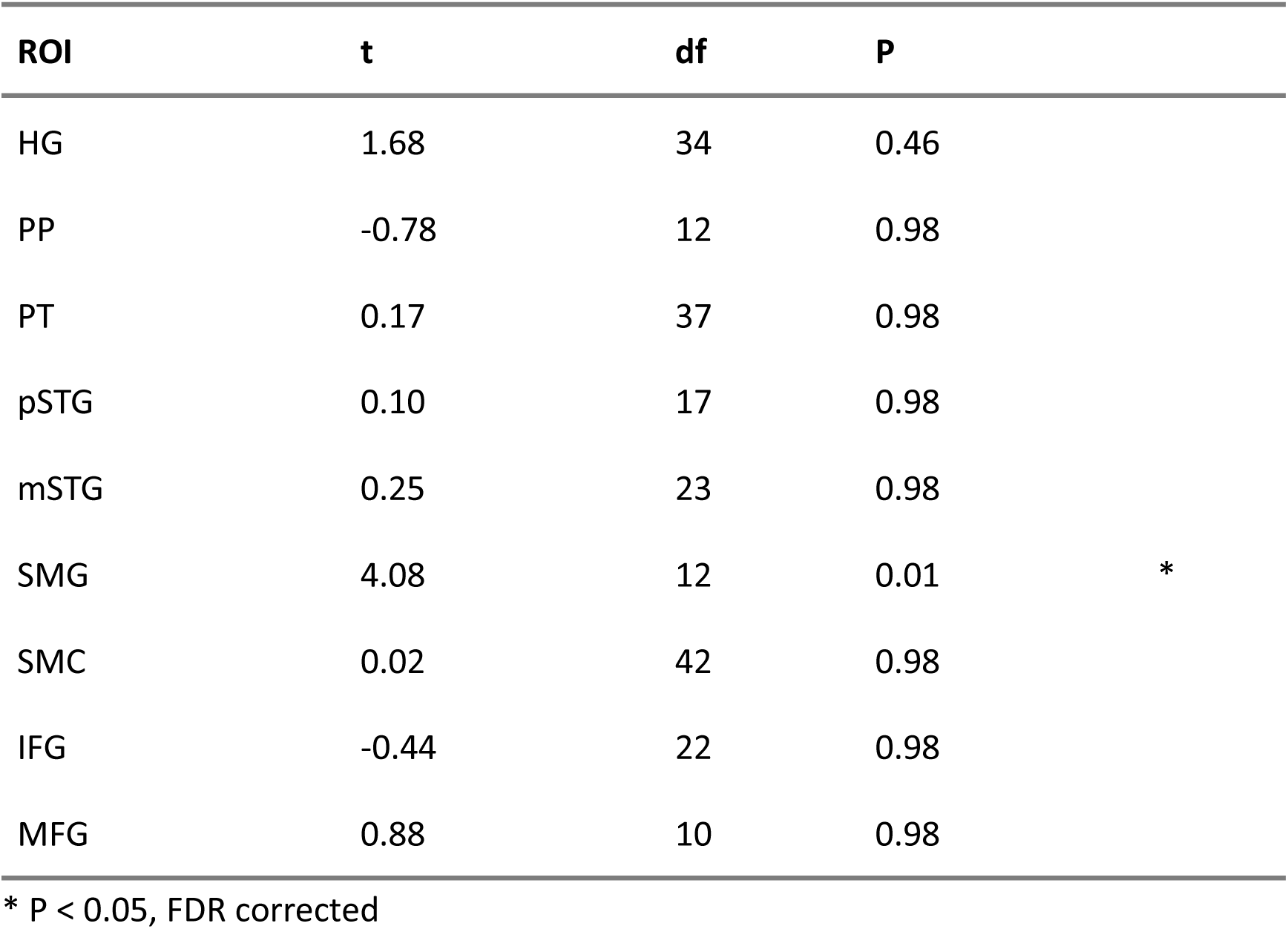
One-sample t-tests, comparing beat index obtained from the autocorrelation of intracerebral EEG responses to the weakly-periodic rhythm to zero.

**S5 Table.** Mean responsiveness z-score in each anatomical region.

| **rhythm** | **ROI** | **responsiveness**<br>**z-score** | **95% low CI** | **95% high CI** |
| --- | --- | --- | --- | --- |
| weakly-periodic | HG | 32.64 | 22.62 | 42.66 |
| weakly-periodic | PP | 7.97 | 5.20 | 10.73 |
| weakly-periodic | PT | 14.59 | 10.59 | 18.59 |
| weakly-periodic | pSTG | 7.71 | 4.92 | 10.51 |
| weakly-periodic | mSTG | 9.68 | 7.11 | 12.25 |
| weakly-periodic | SMG | 8.99 | 5.16 | 12.82 |
| weakly-periodic | SMC | 9.04 | 7.34 | 10.74 |
| weakly-periodic | IFG | 3.56 | 3.07 | 4.05 |
| weakly-periodic | MFG | 2.73 | 2.54 | 2.91 |
| strongly-periodic | HG | 28.64 | 20.29 | 36.99 |
| strongly-periodic | PP | 6.60 | 4.22 | 8.98 |
| strongly-periodic | PT | 12.79 | 10.10 | 15.47 |
| strongly-periodic | pSTG | 6.04 | 4.05 | 8.03 |
| strongly-periodic | mSTG | 8.01 | 5.76 | 10.26 |
| strongly-periodic | SMG | 9.44 | 6.90 | 11.97 |
| strongly-periodic | SMC | 7.53 | 6.02 | 9.04 |
| strongly-periodic | IFG | 2.41 | 1.74 | 3.08 |
| strongly-periodic | MFG | 0.60 | -0.41 | 1.60 |

**S6 Table.**
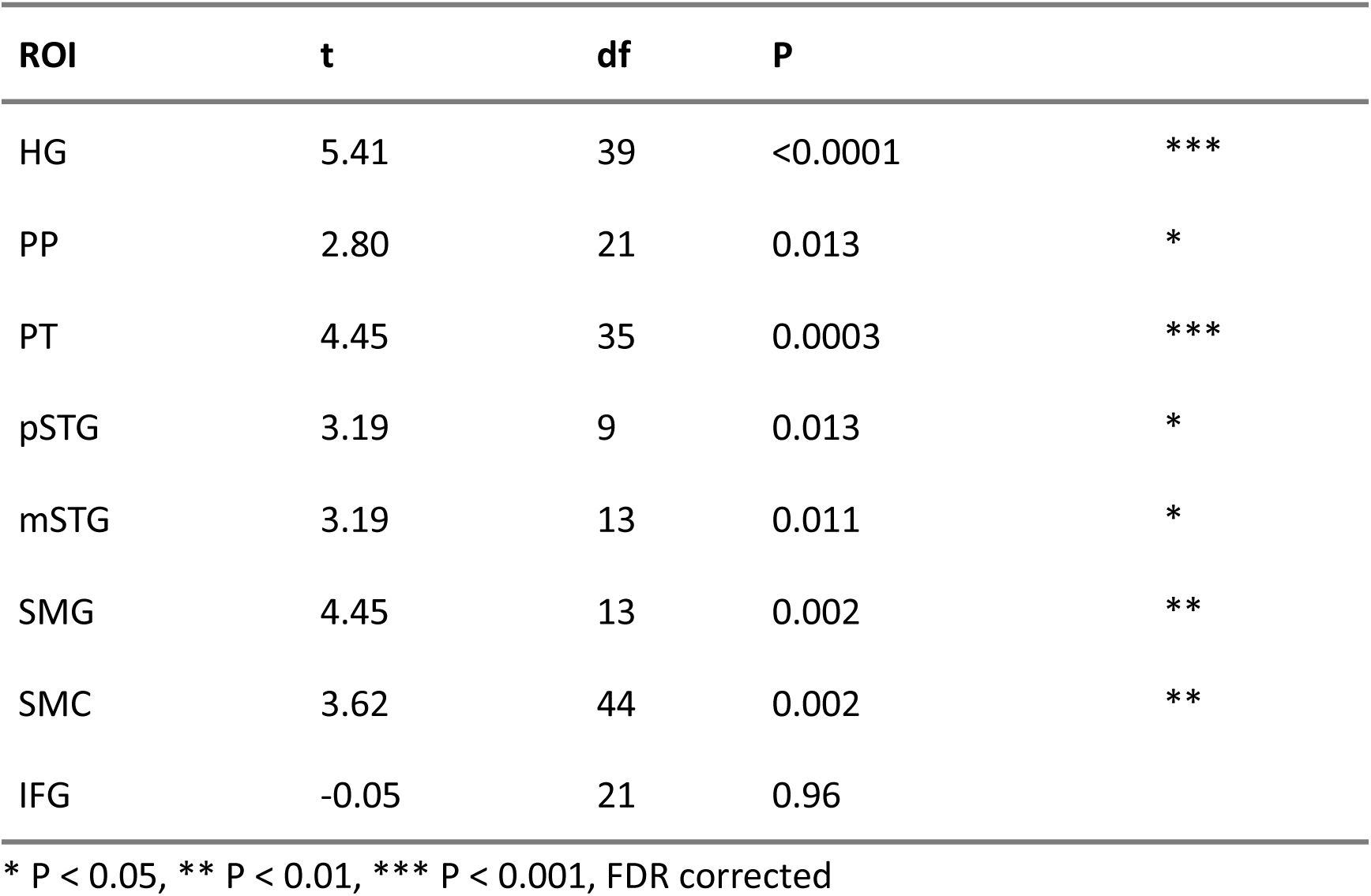
One-sample t-tests, comparing beat index obtained from the magnitude spectrum of bipolar reference intracerebral EEG responses to the weakly-periodic rhythm with the value obtained from the simulated brainstem response.

| ROI | t | df | P |  |
| --- | --- | --- | --- | --- |
| HG | 5.41 | 39 | <0.0001 | *** |
| PP | 2.80 | 21 | 0.013 | * |
| PT | 4.45 | 35 | 0.0003 | *** |
| pSTG | 3.19 | 9 | 0.013 | * |
| mSTG | 3.19 | 13 | 0.011 | * |
| SMG | 4.45 | 13 | 0.002 | ** |
| SMC | 3.62 | 44 | 0.002 | ** |
| IFG | -0.05 | 21 | 0.96 |  |

\* P < 0.05, \*\* P < 0.01, \*\*\* P < 0.001, FDR corrected

**S7 Table.** One-sample t-tests, comparing beat index obtained from the magnitude spectrum of bipolar reference responses to the weakly-periodic rhythm with zero.

| ROI | t | df | P |
| --- | --- | --- | --- |
| HG | 0.17 | 39 | 0.99 |
| PP | -0.50 | 21 | 0.97 |
| PT | 0.00 | 35 | 1 |
| pSTG | 0.90 | 9 | 0.97 |
| mSTG | 0.69 | 13 | 0.97 |
| SMG | 2.37 | 13 | 0.14 |
| SMC | -0.35 | 44 | 0.97 |
| IFG | -2.92 | 21 | 0.07 |

**S1 Fig.**
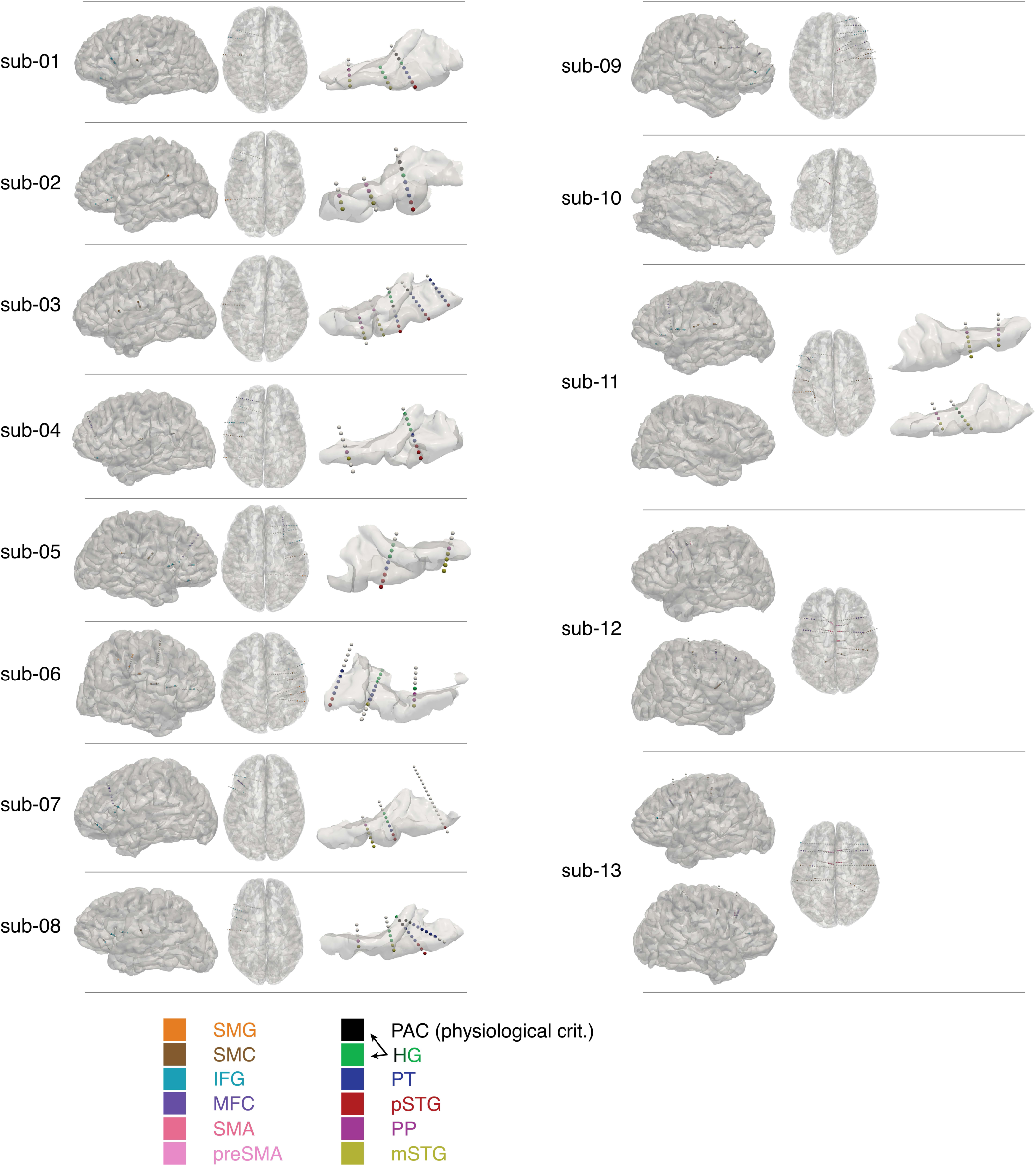
Anatomical distribution of contacts shown on each participant’s pial surface reconstruction. Contacts are colored according to anatomical region. Whole depth electrode is shown if it contained at least one contact localized in a region of interest. Contacts in the auditory regions are shown on the temporal lobe reconstruction. Contacts in the associative regions are shown on the reconstruction of the whole corresponding hemisphere.

**S2 Fig.**
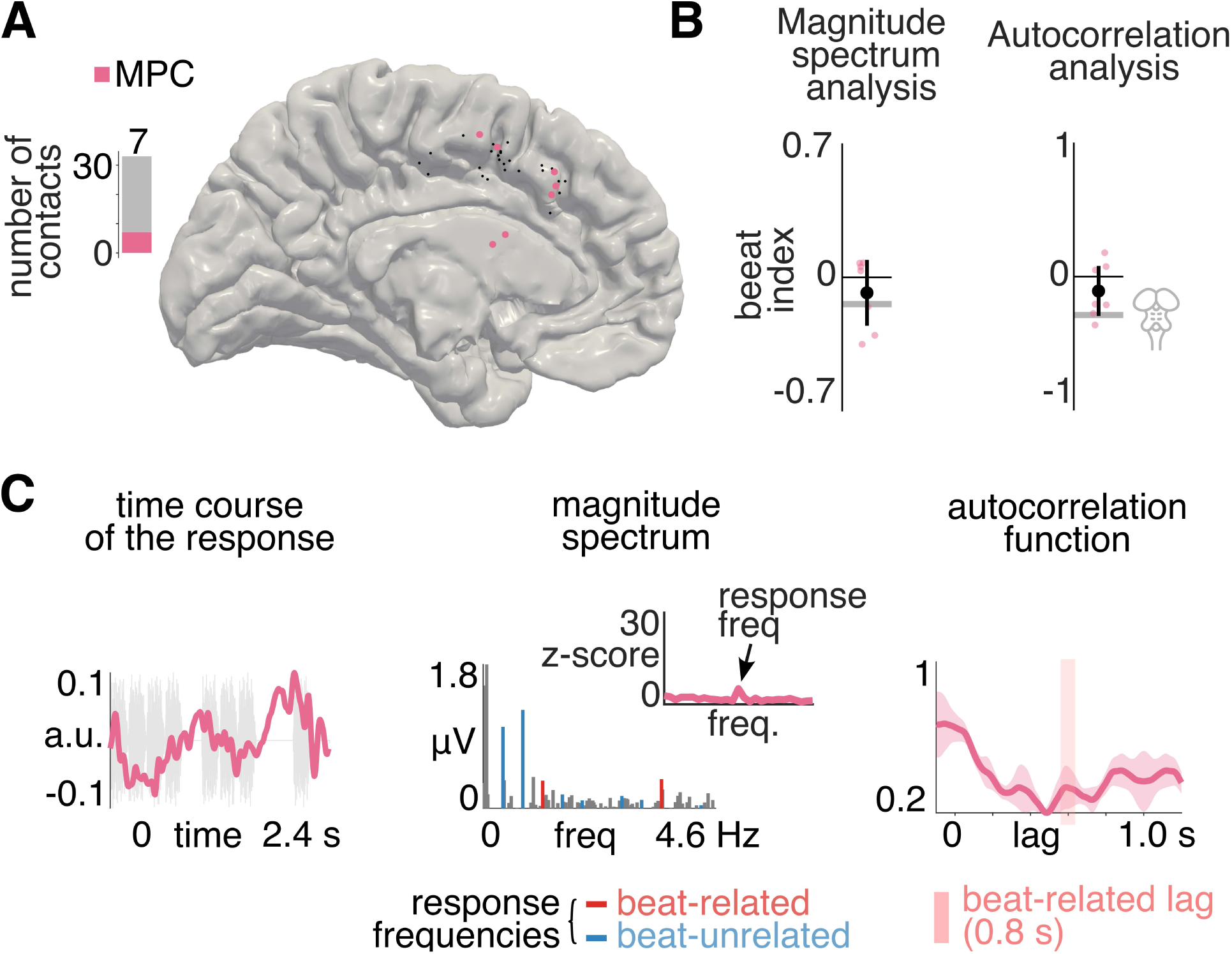
Response to weakly-periodic rhythm in medial premotor cortex (MPC). (A) Spatial distribution of contacts in the MPC projected onto the pial surface of an MNI brain for visualization (collapsed across hemispheres). Note the projection into MNI space may distort the spatial location of individual contacts. Contacts in MPC were labelled as SMA or preSMA based on individual anatomical landmarks and merged into a single region of interest. Each circle represents an individual contact. Responsive contacts are shown in color. (B) Beat index based on magnitude spectrum (left) and autocorrelation (right; see Fig 2C and 2E legend for further details). (C) Summary time course, magnitude spectrum, and autocorrelation function of responsive contacts (see Fig 3 legend for further details).

**S3 Fig.**
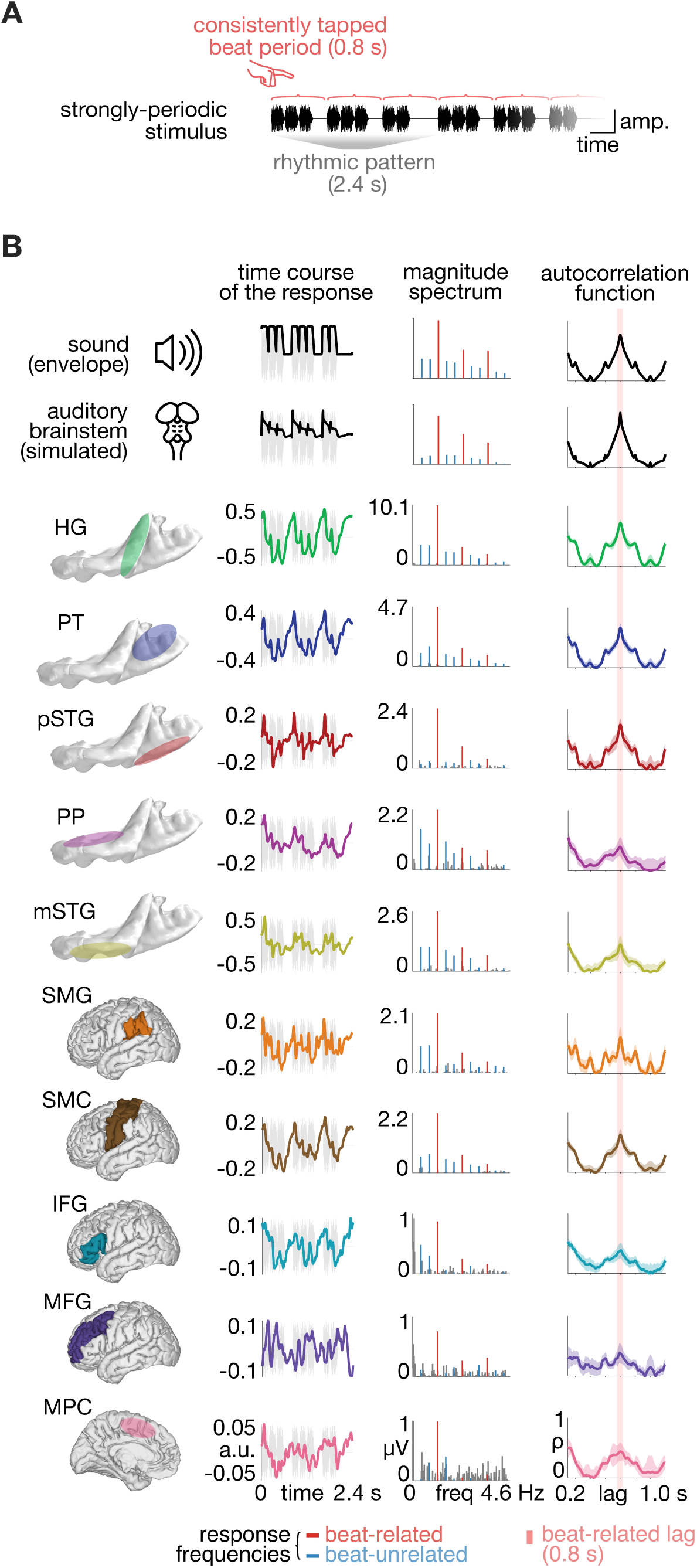
Summary of the response to the strongly-periodic rhythm in each anatomical region. Same as Fig 3, but for responses to the strongly-periodic stimulus sequence.

**S4 Fig.**
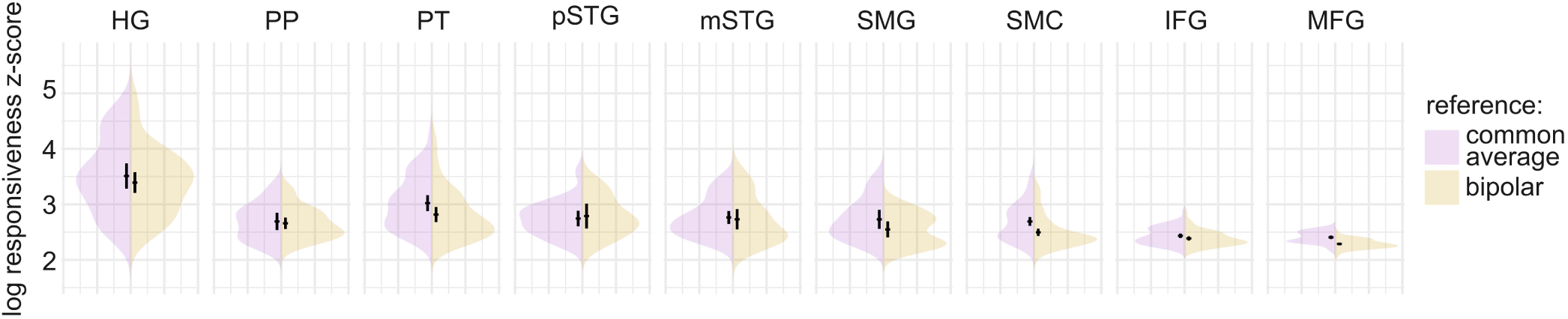
Differences in SNR (log of responsiveness z-score) between the common average and bipolar referencing scheme, separately for each region of interest. The black circles and vertical error bars indicate the mean and 95% CI respectively.

**S1 Text**: Control analysis using bipolar reference montage.

Results reported in the main text were obtained using a common average reference. However, given that different referencing schemes may differentially emphasize or attenuate particular aspects of neural responses [59], we repeated the analyses for the weakly periodic stimulus using a bipolar reference (see Methods).

We used a more lenient threshold to select responsive contacts (1.64, i.e., p < 0.05 one-tailed: signal > noise) as bipolar montage yielded one less channel per depth electrode as well as lower SNR (smaller responsiveness z-score when comparing bipolar channels to common average reference, accounting for the variability across brain regions of interest; linear mixed-effects model, P < 0.001, BF_10_ > 1000, see S4 Fig).

Temporal lobe auditory cortex responses showed overall similar results with bipolar reference as compared to the average reference montage. Specifically, compared to the simulated brainstem response, beat index obtained from the magnitude spectrum was significantly enhanced in the HG, considered the anatomical landmark of the primary auditory cortex (one sample t-test, t_39_ = 5.41, P < 0.0001), and even after applying a strict physiological criterion to isolate the primary auditory cortex (t_17_ = 3.77, P = 0.002). However, neither of these selections revealed a significantly positive beat index (Ps > 0.05). In addition, there was no difference across subfields of the auditory cortex (linear mixed-effects model, P = 0.65, BF_10_ = 0.11), which all showed a significantly greater beat index compared to the subcortical model (Ps < 0.05) but not zero (Ps > 0.05, FDR corrected, see S6 Table).

Regarding responses beyond the temporal lobe auditory cortex, a generally smaller number of responsive contacts was identified across associative regions of interest, even with the more lenient responsiveness criteria, with only 1 responsive contacts in MFG and 2 responsive contacts in SFG, which were therefore both excluded from further analyses. The beat index was significantly different when comparing associative regions and HG in a single linear mixed-effects model (P = 0.002, BF_10_ = 28.42). This was driven by a significantly enhanced beat index in SMG compared to IFG (P = 0.001), SMC (P = 0.03), and marginally HG (p = 0.06). In addition, IFG showed a reduced beat index compared to HG and SMC (P = 0.03 for both, FDR corrected). All other contrasts were nonsignificant. All tested regions except of IFG showed an enhanced beat periodicity compared to the simulated brainstem model (Ps < 0.05, FDR corrected, see S6 Table). While no region showed a significantly positive beat index (Ps > 0.05), a trend towards significance was observed in the SMG (P = 0.07, FDR corrected, see S7 Table). Only 3 responsive contacts were found in the MPC and therefore not further analyzed.

Taken together, while isolating highly local brain activity using a bipolar reference successfully captured periodization between the subcortical and cortical stages of the auditory pathway, this montage was not as sensitive as the common average reference to further transformations in the parietal cortex. This reduction in sensitivity was apparent even with a more lenient responsiveness threshold used here to account for the generally lower SNR yielded by bipolar referencing. It is widely acknowledged that selecting an optimal referencing scheme for capturing specific features of neural activity is a complex issue that depends on multiple factors, including the research question, anatomical location, analysis approach, and the frequency range of interest [59,131,132]. Relevant for the current study, as bipolar montage subtracts activity shared by neighboring electrode pairs, it may be less sensitive to spatially extended fluctuations in field potentials. This limitation has been shown to particularly affect high-amplitude low-frequency activity, possibly due to signal cancellation if the cortical extent of the neural source is equal to or larger to inter-contact distance [59,60]. Given that the present study focused on low-frequency activity (below 5 Hz), which has been identified as critical for beat periodization [8,14,20,61], a global referencing scheme may be better suited to capture the neural transformation processes supporting beat perception.

